# Sex differences in oxycodone-taking behaviors are linked to disruptions in reward-guided, decision-making functions

**DOI:** 10.1101/2024.04.09.587443

**Authors:** Kaitlyn LaRocco, Peroushini Villiamma, Justin Hill, Mara A. Russell, Ralph J. DiLeone, Stephanie M. Groman

## Abstract

Problematic opioid use that emerges in a subset of individuals may be due to pre-existing disruptions in the biobehavioral mechanisms that regulate drug use. The identity of these mechanisms is not known, but emerging evidence suggests that suboptimal decision-making that is observable prior to drug use may contribute to the pathology of addiction and, notably, serve as a powerful phenotype for interrogating biologically based differences in opiate-taking behaviors. The current study investigated the relationship between decision-making phenotypes and opioid-taking behaviors in male and female Long Evans rats. Adaptive decision-making processes were assessed using a probabilistic reversal-learning task and oxycodone- (or vehicle, as a control) taking behaviors assessed for 32 days using a saccharin fading procedure that promoted dynamic intake of oxycodone. Tests of motivation, extinction, and reinstatement were also performed. Computational analyses of decision-making and opioid-taking behaviors revealed that attenuated reward-guided decision-making was associated with greater self-administration of oxycodone and addiction-relevant behaviors. Moreover, pre-existing impairments in reward-guided decision-making observed in female rats was associated with greater oxycodone use and addiction-relevant behaviors when compared to males. These results provide new insights into the biobehavioral mechanisms that regulate opiate-taking behaviors and offer a novel phenotypic approach for interrogating sex differences in addiction susceptibility and opioid use disorders.

## Introduction

The emergence and persistence of problematic opioid-taking behaviors has been proposed to be driven by pre-existing differences in the neurobiological mechanisms that regulate drug use (Evans and Cahill, 2016; Marie and Noble, 2023). The identity of these mechanisms is not known, in part, because dissociating addiction susceptibility mechanisms from the robust drug-induced neural adaptations that occur in response to drug exposure has been a challenge. Emerging evidence suggests that decision-making could be a useful behavioral biomarker for delineating the biological mechanisms that mediate addiction susceptibility from those that are consequent to drug use (Groman and Jentsch, 2012; Groman et al., 2022). Specifically, we have found that deficits in reward-guided, decision-making prior to any drug exposure are predictive of greater cocaine-taking behaviors in rats (Groman et al., 2020; Villiamma et al., 2022) and associated with neurobiological abnormalities that have been observed in cocaine-dependent individuals (Volkow et al., 1993; Groman et al., 2011, 2020; Payer et al., 2014). Decision-making phenotypes could be used for assessing addiction risk in individuals prior to any drug exposure and, therefore, represent a potentially powerful approach for impeding the ongoing opioid epidemic.

The rapid rise in opioid misuse and abuse has been linked to prescription opioids such as oxycodone (Kenan et al., 2012; Evans et al., 2018; Kibaly et al., 2020).

Approximately ∼13% of oxycodone users will abuse the drug and there is evidence that this prevalence is even higher in females compared to males (Han et al., 2017; Richesson and Jennifer M. Hoenig, 2021; Carrasco-Garrido et al., 2022). Moreover, many heroin users report that initiation of opiates began following use of oxycodone (Lankenau et al., 2012; Monico and Mitchell, 2018). Despite the profound risks associated with oxycodone, it remains one of the most widely prescribed painkillers and there is significant interest in understanding the neurobiology that leads to abuse and misuse of oxycodone (Webster, 2017; Moningka et al., 2019; Levis et al., 2021; Oswald et al., 2021). Several new oral self-administration procedures have been developed to interrogate the neurobiology of opioid use disorder in rodents (Slivicki et al., 2023).

Many of these oral procedures lead to patterns of oxycodone use that are different to those observed in human populations, including uniformly high rates of escalation and intake of oxycodone (Enga et al., 2016) in both male and female subjects (Slivicki et al., 2023, however also see Fulenwider et al., 2020). Given that most oxycodone users do not develop abuse the drug (Richesson and Jennifer M. Hoenig, 2021), but that risk for abuse is elevated in women compared to men, developing an oral self-administration procedure that is able to model the biologically-mediated variation in oxycodone use is critical for understanding the neurobiology of opioid use disorder.

The current study sought to understand the relationships between decision-making functions and oxycodone-taking behaviors in both male and female rats. We hypothesized that deficits in reward-guided decision-making prior to any drug exposure would be predictive of greater oxycodone intake and, notably, that these biobehavioral relationships would differ between male and female rats. To test this hypothesis, decision-making functions were assessed in adult, male and female Long Evans rats using a three-choice, spatial probabilistic reversal learning task. Oxycodone-taking behaviors were assessed using a novel, oral self-administration procedure in 3 h daily sessions for 32 days followed by tests of motivation, extinction, and reinstatement. We report that reward-guided decision-making was predictive of subsequent oxycodone self-administration and that greater oxycodone self-administration in female rats was associated with poorer reward-guided, decision-making compared to male rats.

## Materials and Methods

### Subjects

Male (N=78) and female (N=78) Long Evans rats were purchased from Charles River Laboratories at approximately 6 weeks of age. Rats were pair-housed in a climate-controlled vivarium on a 10 h light/dark cycle (lights on at 6am; lights off at 8pm). Rats had ad libitum access to water and underwent dietary restriction to 90% of their free-feeding body weight throughout the experiment. Experimental procedures were approved by the Institutional Animal Care and Use Committee (IACUC) at the University of Minnesota and according to the National Institutes of Health institutional guidelines and Public Health Service Policy on humane care and use of laboratory animals.

### Probabilistic reversal learning (PRL) task

Decision-making was assessed on a three-choice, spatial PRL task using stochastic reward schedules in operant conditioning chambers (Med Associates; Figure 1A). Rats initiated trials by making a noseport entry into a magazine. This led to the illumination of three noseport apertures on the opposite panel. One of these apertures was associated with a higher probability of delivering reward than the other two apertures (Figure 1B). Reinforcement probabilities assigned to each noseport were pseudorandomly assigned at the start of each session for individual rats. Rats could make a single choice on each trial by making a noseport entry into the illuminated port. Rats were initially trained on a dynamic, variable schedule of reinforcement in which the reinforcement probabilities assigned to each noseport changed based on the performance of the rat (referred to as the *variable PRL schedule;* Figure 1B). For example, at the start of each session each noseport was pseudo-randomly assigned to deliver reward with a probability of 70, 30, or 10% by the program. When rats met a performance criterion (21 choices on the highest reinforced noseport in the last 30 trials), the probabilities reversed between two noseports: the highest reinforced noseport (70%) became the lowest reinforced noseport (10%) whereas the lowest reinforced noseport (10%) became the highest reinforced noseport (70%). The noseport associated with a probability of 30% reinforcement remained unaltered. These reinforcement probabilities remained unchanged until the performance criterion was once again met, after which the reinforcement probabilities reversed again between the noseports associated with the highest and lowest reinforcement probabilities. The occurrence of a reversal was contingent upon performance and rats could complete as many as 8 reversals in a single session. Sessions terminated when rats completed 250 trials or 75 min had lapsed, whichever occurred first.

**Figure 1:**
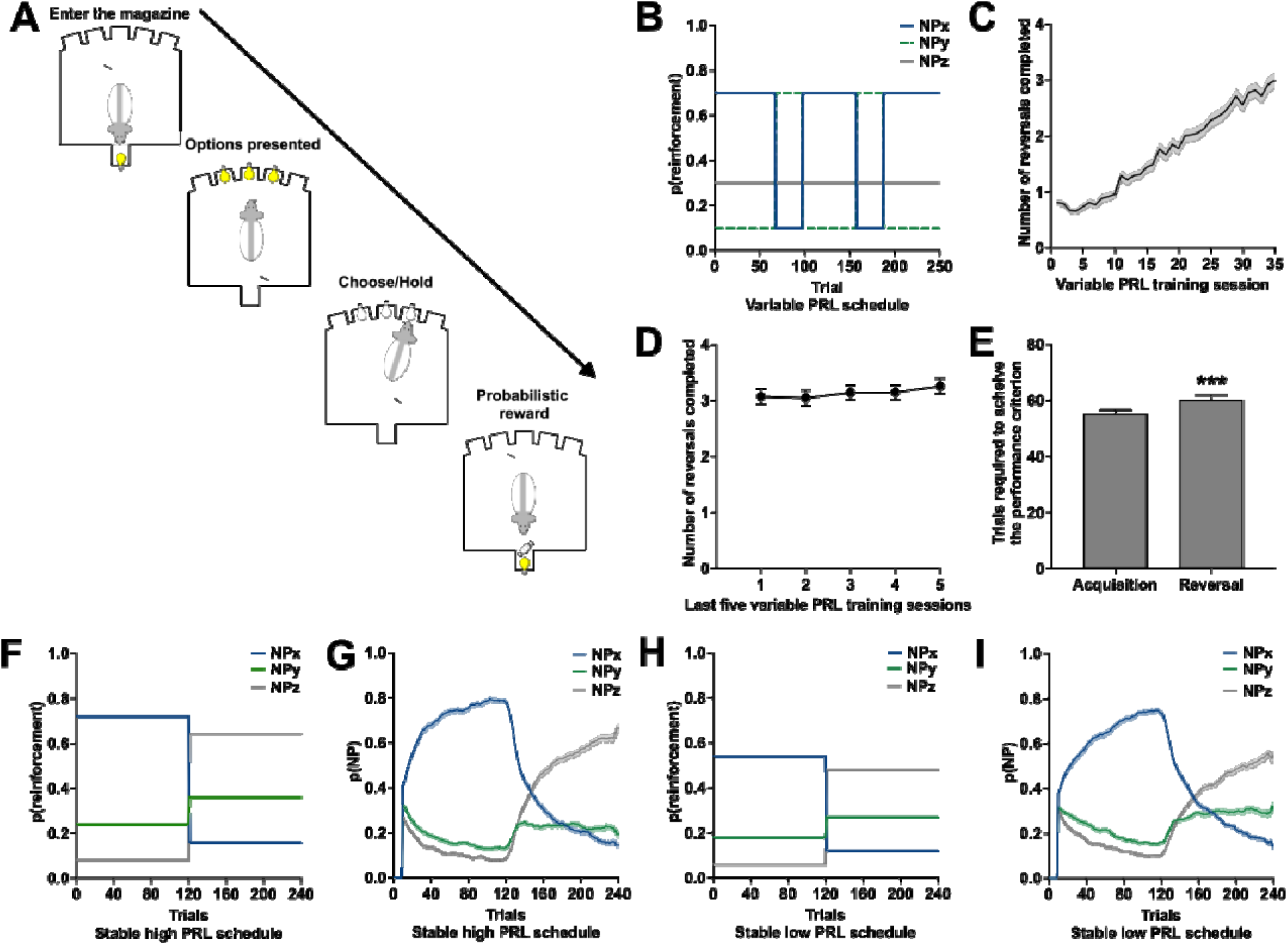
The probabilistic reversal learning (PRL) task. (A) Schematic of a single trial event. (B) The probability of reinforcement for the three noseport apertures under the variable schedule of reinforcement. (C) The number of reversals rats were able to complete on the variable schedule of reinforcement increased across the 35 training sessions. (D) The number of reversals rats completed in the last five sessions on the variable schedule of reinforcement was stable. (E) The number of trials rats required to acquire a discrimination was less than that required to reverse the discrimination. (F) The probability of reinforcement for the three noseport apertures under the stable high schedule of reinforcement. (G) The choice behavior of rats on the stable high schedule of reinforcement. (H) The probability of reinforcement for the three noseport apertures under the stable low schedule of reinforcement. (G) The choice behavior of rats on the stable low schedule of reinforcement.

Rats completed 35 sessions on the variable PRL schedule and then the stability in the number of reversals that rats completed in a single session was assessed. If the standard deviation in the number of reversals across the last five sessions was greater than 1, then rats completed additional variable PRL sessions until this stability criterion was met or rats had completed a total of 50 variable PRL sessions, whichever occurred first. Decision-making functions were also assessed using two other schedules of reinforcement where a single reversal occurred when rats had completed 120 trials (Figure 1F, 1H). The *stable high PRL schedule* had a larger dynamic range compared to the *stable low PRL schedule* (Figure 1F, 1H). Sessions terminated when rats completed 240 trials or 75 min had lapsed, whichever occurred first.

Performance in the PRL task across the three schedules of reinforcement was highly correlated with one another (Supplemental Figure 1), so dependent measures were collapsed across the different schedules of reinforcement to limit the number of comparisons made in the current study and improve the accuracy of the decision-making estimates.

### Oral self-administration paradigm

Following the decision-making assessments in the PRL task, rats were divided into two groups based on their performance in the PRL task: rats were trained to self-administer either a vehicle solution (0.05% saccharin in water; N=41) or oxycodone solution (0.05 mg/kg/infusion of oxycodone in 0.05% saccharin solution; N=115) in 3 h daily sessions for 32 days in a novel operant environment (Figure 2A). The self-administration procedure is described in detail in the Supplement. Briefly, entries into the active noseport resulted in delivery of the vehicle or oxycodone solution directly into the magazine (∼150 ul of solution). Rats were trained on a fixed-ratio (FR) 1 schedule for three days to establish operant responding. The operant requirement was then increased to a FR3 schedule for the remaining self-administration sessions. Responses in the inactive noseport were recorded but had no programmed consequence. On the 9^th^ self-administration session, the dose of oxycodone was doubled (0.05 mg/kg/infusion to 0.1 mg/kg/infusion) and the saccharin gradually removed from the vehicle solution (Figure 2B) such that water was the vehicle solution for both groups by the thirteenth self-administration session.

**Figure 2:**
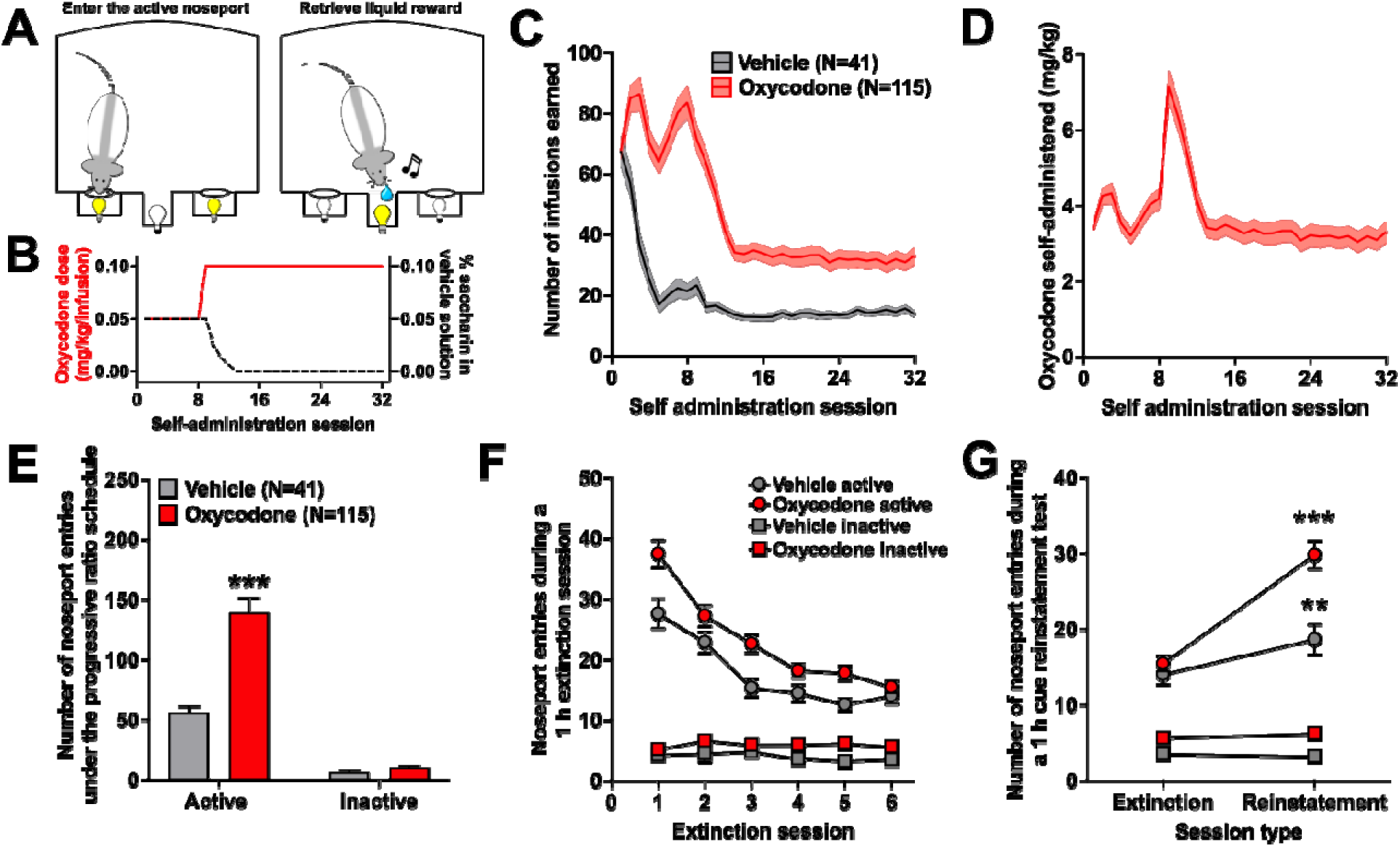
Oral self-administration of oxycodone in rats. (A) Schematic of the operant environment. (B) Dose of oxycodone and percentage of saccharin in the vehicle solution across the 32 self-administration sessions. (C) The total number of infusions rats in the vehicle (N=41) and oxycodone (N=115) earned across the 32 self-administration sessions. (D) The amount of oxycodone consumed (mg/kg) across the self-administration sessions. (E) Number of active and inactive responses rats made under the progressive ratio schedule of reinforcement in the vehicle group (gray bars) and oxycodone group (red bars). (F) The number of active (circles) and inactive (squares) noseport responses rats in the vehicle (gray bars) and oxycodone (red bars) groups made across the six 1 h extinction sessions. (G) The number of active (circles) and inactive (square) noseport responses rats in the vehicle (gray bars) and oxycodone (red bars) groups made in the 1 h cue-induced reinstatement test. ** p<0.01; *** p<0.001.

### Addiction-relevant behaviors

Additional tests of drug–seeking and –taking behaviors were conducted following the self-administration sessions using procedures previously described (Groman et al., 2020). Motivation to obtain an infusion of oxycodone or vehicle was assessed under a progressive schedule, the ability to inhibit non-reinforced, drug-taking behaviors was assessed under extinction, and drug-seeking behavior assessed in a cue-induced reinstatement test. Details of these procedures are described in the Supplement.

## Data analysis

### Reinforcement-learning model

Choice behavior collected from rats in the PRL task was analyzed with a differential forgetting reinforcement-learning model (Barraclough et al., 2004; Ito and Doya, 2009; Groman et al., 2019a), which captures gradually decaying effects of previous choices and outcomes on choices. This model was found to fit the choice behavior of rats significantly better than other reinforcement-learning models (see Supplemental methods and Supplemental Table 1), as we have previously shown (Groman et al., 2020; Moin Afshar et al., 2022; Villiamma et al., 2022). This reinforcement-learning model contains four free parameters: a decay rate for the action values of chosen options (γ_C_), a decay rate for the action values of unchosen options (γ_U_), a parameter for the appetitive strength of rewarded outcomes (Δ_+_), and a parameter for the aversive strength of unrewarded outcomes (Δ_0_).

### Characterizing latent drug-taking phenotypes

The number of oxycodone infusions that individual rats self-administered in the last 20 self-administration sessions, when the vehicle no longer contained saccharin, were fit with a power function:

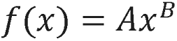

where X is the session number and A and B are free parameters estimating the scaling factor and the rate of growth, respectively. The A parameter determines the initial strength of drug-taking behaviors: rats with a higher A parameter self-administered more oxycodone than rats with a lower A parameter across the self-administration procedure (Supplemental Figure S3). The B parameter determines the growth of the function: rats with a higher, positive B parameter escalated in the number of oxycodone infusions they self-administered compared to rats with a lower, negative B parameter (Supplemental Figure S3).

### Statistical analyses

Values reported are the mean ± SEM, unless otherwise noted. Analyses were performed in SPSS (version 29; IBM Corp., Armonk NY). Repeated measures data were analyzed using a generalized estimating equations (GEE) model using a probability distribution based on the known properties of the data. Specifically, event data (e.g., number of correct choices) were analyzed using a binary logistic distribution and count data (e.g., number of infusions) were analyzed using a negative binomial distribution. Between subjects factors included in the GEE model could be experimental group (oxycodone vs. vehicle) and/or sex (male vs. female). Relationships between dependent measures (e.g., reinforcement-learning parameters and drug-taking behaviors) were analyzed using linear regression models. Mediation analyses were conducted using multiple regression analyses. The indirect effect was calculated using the PROCESS macro in SPSS (Hayes, 2018).

## Results

### Decision-making is reliable across different schedules of reinforcement

Long Evans rats (male N=78; female N=78) were trained to make flexible choices in a three-armed bandit probabilistic reversal-learning (PRL) task (Figure 1A) using stochastic reward schedules (Figure 1B). The number of reversals that rats completed in a single session improved across the 35 training sessions under the variable schedule of reinforcement (Figure 1C; session: Wald x^2^=825; p<0.001) and stabilized after completing 36.23 ± 0.49 sessions (Figure 1D; Cronbach’s a=0.86). The number of trials that rats required to reach the first performance criterion was significantly less than that required to achieve the second performance criterion (Figure 1E; Wald x^2^=1069; p<0.001) indicating that despite the extensive training rats received on the PRL task they still found it difficult to reverse a discrimination.

Choice behavior under the stable high and stable low schedules of reinforcement was then examined (Figure 1F-K). Rats chose the most frequently reinforced option both before and after the reversal at a rate significantly greater than chance (all t(151)>8.50; all p’s<0.001; Supplemental Figure 2). Moreover, the probability of choosing the highest reinforced option following the reversal was lower than pre-reversal performance (Wald x^2^=485; p<0.001) and performance under the stable low schedule of reinforcement was lower than that under the stable high schedule (Wald x^2^=54.06; p<0.001). Rats were, therefore, able to track dynamic, as well as novel, schedules of reinforcement.

Choice behavior of rats were fitted with the four different computational algorithms (see description above and in the Supplement) and the Bayesian Information Criterion (BIC) calculated for each model (Table 1). The differential forgetting reinforcement learning model had the lowest BIC value and, therefore, best fit the choice data collected in rats. The average parameter estimates obtained from the differential forgetting reinforcement learning model are presented in Table 2.

**Table 1:** The total bayesian information criterion (BIC) for each of the four computational algorithms. The differential forgetting reinforcement learning model (Model 4) had the lowest BIC value compared to all other models, indicating that this model best fit the rat choice data.

|  | Model 1: Q learning | Model 2: Forgetting Q learning | Model 3: Forgetting reinforcement learning | Model 4: Differential forgetting reinforcement learning |
| --- | --- | --- | --- | --- |
| BIC | 3490441 | 3159931 | 3034649 | **2995183** |

**Table 2:** Quartile values of the parameter estimates from the DF reinforcement learning mode.

| | $\gamma_{\mathbf{c}}$ | $\gamma_{\mathbf{u}}$ | $\Delta_{+}$ | $\Delta_{\mathbf{0}}$ |
| --- | --- | --- | --- | --- |
| 25<sup>th</sup> | 0.59 | 0.67 | 1.08 | -0.02 |
| Median | 0.65 | 0.79 | 1.59 | 0.21 |
| 75<sup>th</sup> | 0.74 | 0.89 | 2.12 | 0.43 |

### Oral self-administration of oxycodone is reinforcing and leads to addiction-relevant behaviors

Following the decision-making assessments, rats were trained to self-administer an oral solution of either oxycodone (0.05 mg/kg/infusion in 0.05% saccharin solution; N=115) or vehicle (0.05% saccharin solution; N=41) in 3 h daily sessions for 32 days (Figure 2A). The dose of oxycodone was doubled on the ninth self-administration session (0.1 mg/kg/infusion) and saccharin gradually eliminated from the vehicle solution (Figure 2B) so that no saccharin was present in the oxycodone or vehicle solution on the 13^th^ self-administration session. The number of infusions rats earned in each of the self-administration sessions is presented in Figure 2C and the amount of oxycodone consumed (mg/kg) presented in Figure 2D.

The number of infusions rapidly decreased in both the vehicle (Wald x^2^=127.98; p<0.001; Figure 2C) and the oxycodone groups (Wald x^2^=51.97; p<0.001; Figure 2C) within the first twelve self-administration sessions. This was likely due to the change in reinforcement schedule (e.g., from FR1 to FR3) and removal of saccharin from the vehicle solution that occurred during this period of operant training. The number of infusions rats earned from the thirteenth self-administration session to the last self-administration session, however, did not significantly change in either the vehicle (Wald x^2^=1.64; p=0.20; Figure 2C) or oxycodone groups (Wald x^2^=0.87; p=0.35). Notably, the number of infusions self-administered by the oxycodone group was significantly greater than the vehicle group (Wald ^2^=56.86; p<0.001) indicating that rats found the oral oxycodone solution to be reinforcing.

Addiction-relevant behaviors were then assessed: oxycodone-taking behaviors were assessed in a single session under a progressive ratio schedule of reinforcement, six days of extinction training, and a cue-induced reinstatement test. The oxycodone group, compared to the vehicle group, made significantly more active responses under the progressive ratio schedule of reinforcement (Wald ^2^=50.75; p<0.001; Figure 2E), during the extinction sessions (Wald ^2^=9.48; p=0.002; Figure 2F), and in the cue-induced reinstatement test (Wald ^2^=71.94; p<0.001; Figure 2G). These data indicate that our oral oxycodone self-administration paradigm led to many of the problematic drug-seeking and -taking behaviors that are observed in opioid-dependent individuals.

### Quantifying individual differences in oxycodone-taking behaviors

The amount of oxycodone that rats self-administered did not increase across the saccharin-free self-administration sessions (session: Wald x^2^=0.87; p=0.35; Figure 2D). This was surprising given recent studies reporting high rates of escalation in oral oxycodone intake in mice (Slivicki et al., 2023) as well as in rats (Zanni et al., 2020). We did, however, observe robust variation in the patterns of oxycodone intake across individual rats and characterized these differences using a power function (Figure 3A), as we have previously done (Groman et al., 2019b, 2020). The distribution of the A and B parameters are presented in Figure 3B and 3C, respectively, and a visualization of how these parameters capture differences in oxycodone use presented in Supplemental Figure 3. The A and B parameters explained nonoverlapping portions of variance in the total amount of oxycodone consumed (R^2^=0.92; A parameter: /J=0.93; p<0.001; B parameter: /J=0.23; p<0.001) and, notably, were not related to one another (R^2^=0.0002; p=0.88; Figure 3D) indicating that these parameter estimates were explaining distinct and unique patterns of oxycodone use. These data highlight the utility of the power function for characterizing non-overlapping, latent oxycodone-taking behaviors.

**Figure 3:**
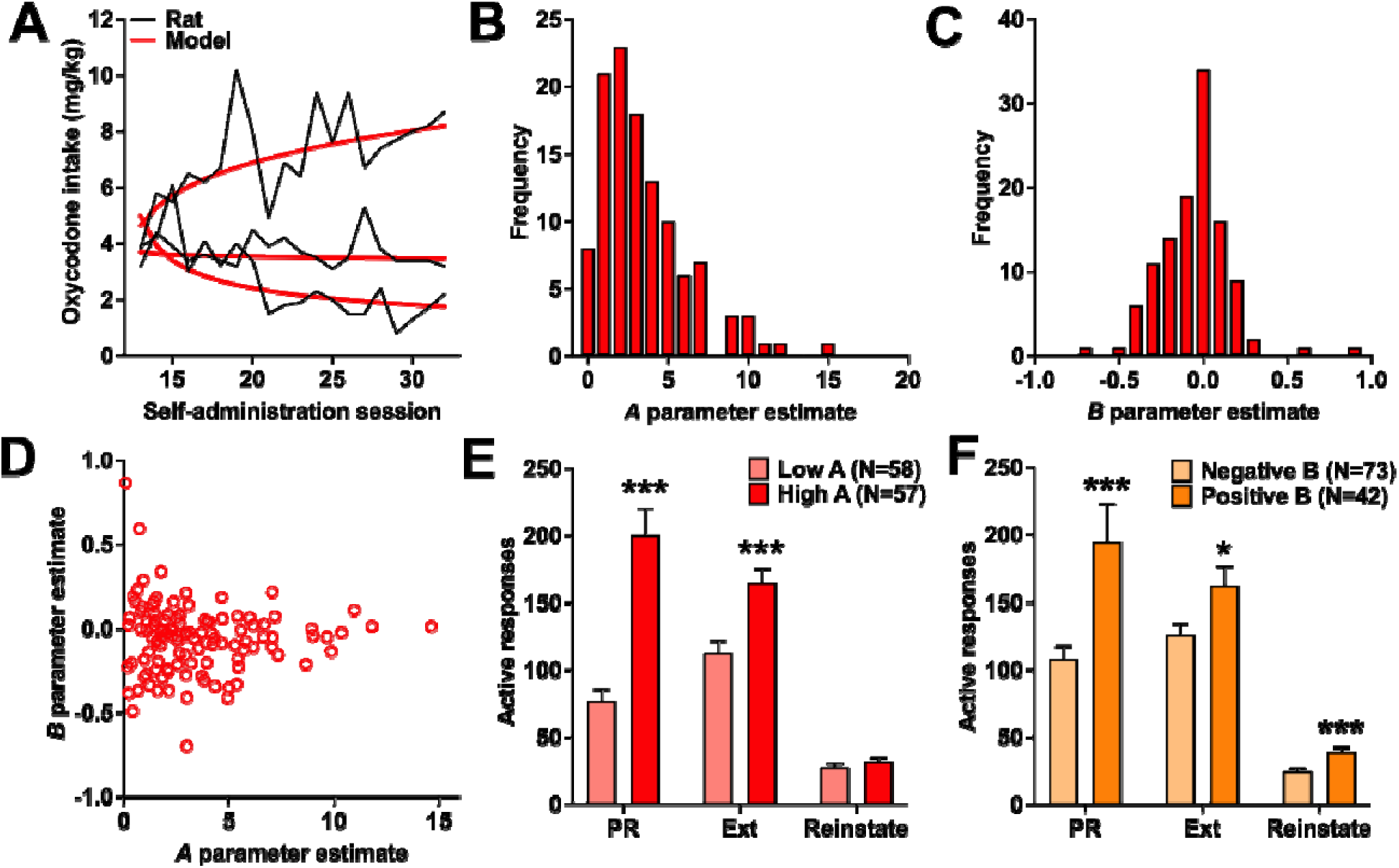
Characterizing latent oxycodone-taking behaviors with the power function. (A) The power function (red lines) was able to capture latent drug-taking behaviors. (B) The frequency distribution of the A parameter and the (C) B parameter estimate. (D) No relationship between the A parameter and B parameter were detected. (E) Active noseport responses in rats with a low A parameter (N=58) and a high A parameter (N=57) in the progressive ratio (PR) test, extinction, and reinstatement. (F) Active noseport responses in rats with a positive B parameter (N=42) and a negative B parameter (N=43) in the progressive ratio (PR) test, extinction, and reinstatement. * p<0.05; *** p<0.001.

Variation in these latent drug-taking behaviors may differentially impact the development of addiction-relevant behaviors. The A and B parameter estimates, but not the interaction between the two parameters, explained a significant amount of variance in responding under the progressive ratio schedule of reinforcement (A parameter Wald ^2^=109.21; p<0.001; B parameter Wald ^2^=47.51; p<0.001; A parameter x B parameter Wald ^2^=1.57; p=0.21), extinction (A parameter Wald ^2^=28.26; p<0.001; B parameter Wald ^2^=23.52; p<0.001; A parameter x B parameter Wald ^2^=0.39; p=0.53), and in the reinstatement test (A parameter Wald ^2^=8.41; p=0.004; B parameter Wald ^2^=37.37; p<0.001; A parameter x B parameter Wald ^2^=0.92; p=0.34; Figure 3E,F). These data suggest that differences in both the A and B parameter are contributing to the emergence of these addiction-relevant behaviors.

### Poor reward-guided, decision-making is predictive of greater oxycodone use

The relationship between decision-making processes that were assessed prior to any drug exposure and oxycodone self-administration was then examined. Rats were divided into two groups based on a median split of performance in the PRL task into rats with good PRL performance (referred to as good PRL; n=57) and rats with poorer PRL performance (referred to as poor PRL; n=58; Figure 4A). The number of oxycodone infusions decreased across the self-administration sessions (day: Wald x^2^=155; p<0.001), but the rate of decrease was significantly lower in the poor PRL group (group x day Wald x^2^=4.44; p=0.04; Figure 4B). Rats in the poor PRL group took significantly more oxycodone than rats in the good PRL group (Wald x^2^=11.05; p<0.001; Figure 4C). The relationship between decision-making processes and self-administration behavior was not observed in the vehicle group (group: Wald x^2^=3.06; p=0.22; group x day: Wald x^2^=3.20; p=0.20; Figure 4D) suggesting that poor decision-making that is predictive of self-administration behavior is specific to oxycodone.

**Figure 4:**
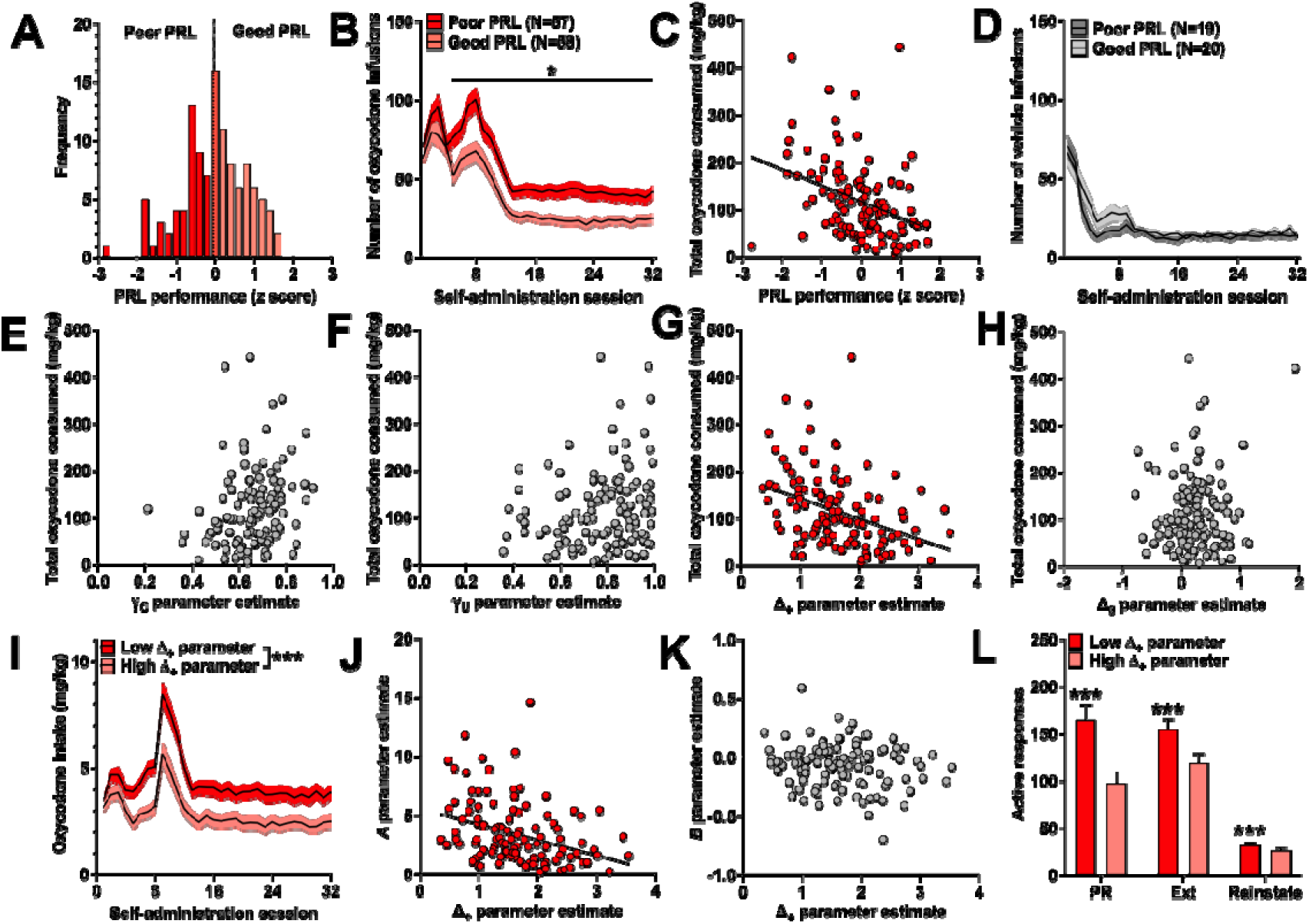
Poor reward-guided decision-making is predictive of problematic oxycodone use. (A) Frequency distribution of performance in the probabilistic reversal learning (PRL) task. Rats were divided into two groups based on a median split of PRL performance: poor PRL (N=57) and good PRL (N=58). (B) Rats in the poor PRL group earned significantly more oxycodone infusions than rats in the Good PRL group. * p<0.05 for group level comparisons. (C) Scatter plot of the relationship between PRL performance and total oxycodone consumed (mg/kg). (D) The number of vehicle infusions rats earned did not differ between poor and good PRL performance. (E) Individual differences in the C were not predictive of oxycodone self-administration. (F) Individual differences in the U parameter were not predictive of oxycodone self-administration. (G) Individual differences in the Δ+ parameter were predictive of oxycodone self-administration. (H) Individual differences in the Δ0 parameter were not predictive of oxycodone self-administration. (I) Rats with a low Δ+ parameter self-administered greater amounts of oxycodone compared to rats with a high Δ+ parameter. (J) Variation in the Δ+ parameter was correlated the initial strength of oxycodone reinforcement, or the A parameter. (K) Variation in the Δ+ parameter was not correlated the rate of growth in oxycodone use, or the B parameter. (L) Rats with a low Δ+ parameter made more active response in the progressive ratio (PR) test, extinction (Ext) sessions, and in the cue-induced reinstatement test compared to rats with a high Δ+ parameter. * p<0.05; ** p<0.01; *** p<0.001.

Differences in decision-making that are predictive of oxycodone use may be associated with a specific reinforcement learning mechanism. The four parameter estimates obtained from the differential forgetting reinforcement learning algorithm were entered into a stepwise regression model predicting the total amount of oxycodone consumed by individual rats. The Δ_+_parameter, but not the y_C_, y_U_, or the Δ_0_ parameter, explained a significant amount of variation in the total amount of oxycodone consumed (Figure 4E-H; R^2^=0.14; p<0.001). Rats with a low Δ_+_parameter took significantly more oxycodone than rats with a high Δ_+_ parameter across the self-administration paradigm (Figure 4I; Wald x^2^=561; p<0.001). Indeed, the Δ_+_ parameter explained a significant amount of variance in the initial strength of oxycodone reinforcement (e.g., A parameter; R^2^=0.11; p<0.001; Figure 4J), but not the rate of change in oxycodone use (e.g., B parameter; R^2^=0.02; p=0.11; Figure 4K).

Disruptions in reward-guided decision-making prior to drug exposure may also facilitate the development of addiction-relevant behaviors. The Δ_+_ parameter was included as a factor in models predicting active lever responses under the progressive ratio schedule of reinforcement, extinction, or the cue-induced reinstatement test. The Δ_+_ parameter explained a significant amount of variance in the number of active responses made under the progressive ratio schedule (Wald x^2^=1306; p<0.001), during the extinction sessions (Wald x^2^=366; p<0.001), and in the cue-induced reinstate test (Wald x^2^=32.41; p<0.001). Rats with a lower Δ_+_ parameter made more active responses compared to rats with a higher Δ_+_ parameter (progressive ratio: /J= -0.46; extinction: -0.23; reinstatement: -0.14; Figure 4H). These data, collectively, indicate that poor reward-guided, decision-making is associated with heightened oxycodone reinforcement that drives elevated intake of oxycodone and heightened addiction-relevant behaviors.

### Oxycodone self-administration is greater in female rats compared to male rats

Emerging evidence indicates that females are be more likely than males to abuse oxycodone. To investigate whether sex differences were present in the current study, self-administration behaviors were compared between male and female rats in the oxycodone and vehicle groups. Post hoc analyses of the significant sex x day interaction (Wald x^2^=41.34; p<0.001) indicated that female rats earned a greater number of oxycodone infusions (Figure 5A) and a greater amount of oxycodone (Figure 5B) compared to male rats. Notably, differences between males and females in oxycodone intake emerged when saccharin had been removed from the vehicle solution and were not observed in the vehicle group (main effect of sex: Wald x^2^=2.74; p=0.10; sex x day interaction: Wald x^2^=2.25; p=0.13; Figure 5C). The parameter estimates characterizing oxycodone-taking behaviors obtained from the power function were then compared between male and female rats. Both the A parameter and B parameter were greater in female rats compared to males (A parameter: Wald x^2^=4.40, p=0.04, /J=1.07; B parameter: Wald x^2^=13.61, p<0.001, /J=0.12) indicating that the initial strength of oxycodone reinforcement (e.g., A parameter) and the rate of growth in oxycodone use (e.g., B parameter) was greater in female rats compared to their male counterparts.

**Figure 5:**
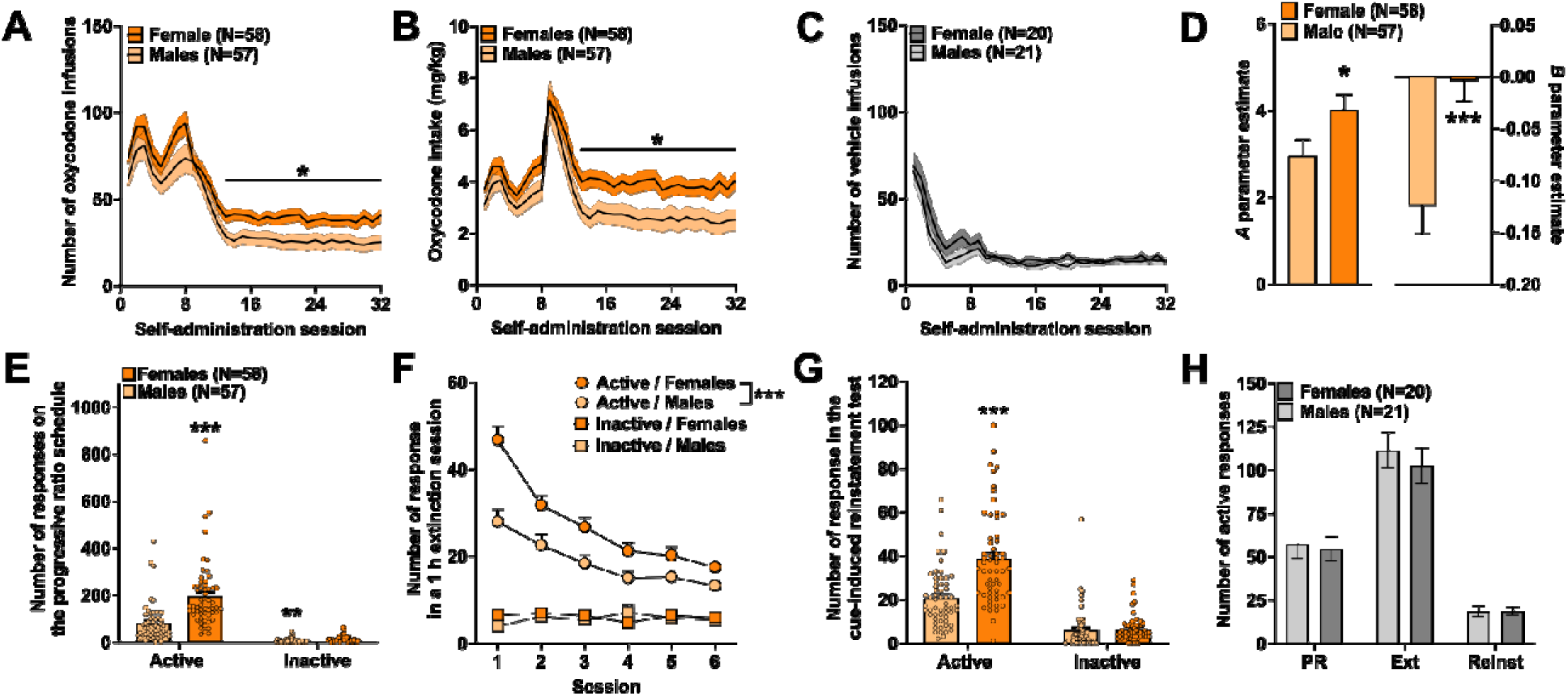
Sex differences in oxycodone-taking behaviors. (A) Female rats earned a greater number of oxycodone infusions compared to male rats. (B) Female rats consumed a greater amount of oxycodone compared to male. (C) The number of vehicle infusions did not differ between males and females. (D) The A parameter and B parameter were larger in female compared to male rats. (E) Females made more responses under the progressive ratio schedule of reinforcement than males. (F) Active responses was greater in female rats compared to males across the six 1 h extinction sessions. (G) Active responses in the cue-induced reinstatement test was greater in females compared to males. (H) No sex differences observed in the vehicle group.

Addiction-relevant behaviors were then compared between male and female rats. Although the number of active and inactive responses made under the progressive ratio schedule of reinforcement was greater in female rats compared to male rats (active responses: Wald x^2^=28.91, p<0.001; inactive responses: Wald x^2^=9.51, p=0.002) the difference between the response types was greater in females (sex x response type:

Wald x^2^=17.45, p<0.001). This indicates that motivation to obtain an oxycodone reinforcer was greater in females compared to males. Moreover, female rats made more active responses across the six extinction sessions (Wald ^2^=16.18, p<0.001; Figure 5F) and during the cue-induced reinstatement test compared to males (Wald ^2^=31.73, p<0.001; Figure 5G) indicating a greater pervasiveness of addiction-relevant behaviors in female rats. These sex differences in addiction-relevant behaviors were specific to the oxycodone group as no sex differences were observed in the vehicle group (Figure 5H; all p’s>0.70).

### Deficits in reward-guided, decision-making in female rats

The results presented here indicate that oxycodone-taking behaviors are elevated in individuals with poor reward-guided, decision-making and, specifically, in female rats compared to male rats. We hypothesized that greater drug-taking behaviors in female rats may be the result of pre-existing disruptions in reward-guided decision-making. Female rats completed a fewer number of reversals compared to male rats across the 35 training sessions under the variable schedule of reinforcement (sex: Wald x^2^=8.87; p=0.003; session: Wald x^2^=596; p<0.001; sex by session interaction: Wald x^2^=2.27; p=0.13; Figure 6A). These differences in the number reversals completed was not driven by differences in the number of trials that female and male rats completed, as sex remained a significant factor when the number of trials was included in the statistical model (Wald x^2^=7.18; p=0.007). Moreover, these sex differences were also observed after performance under the variable schedule of reinforcement had stabilized (Wald x^2^=22.78; p<0.001; Figure 6B) suggesting that these sex differences in PRL performance were not because female rats required additional training sessions to reach the same level of stability as male rats. Similar sex differences were also observed under the other schedules of reinforcement (Supplemental Figure 4). These data, collectively, indicate that female rats had greater difficulty allocating their choices to the highest reinforced option compared to male rats.

**Figure 6:**
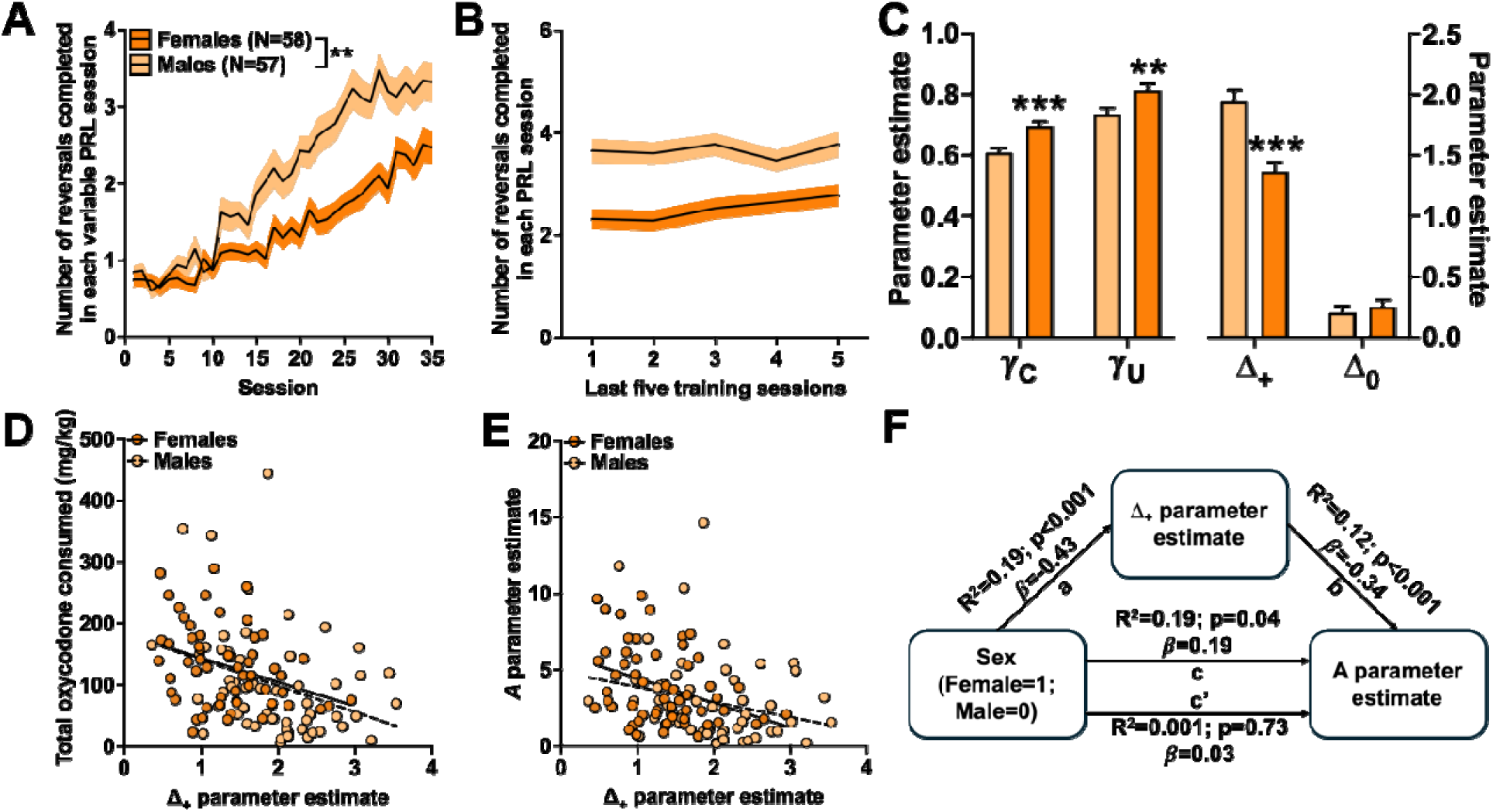
Sex differences in decision-making processes. (A) The number of reversals female (dark orange) and male (light orange) rats completed across the 35 PRL sessions under the variable schedule of reinforcement. (B) The number of reversals female (dark orange) and male (light orange) rats completed once performance had stabilized or rats had completed 50 sessions on the variable PRL task. (C) The average C, U, Δ+, and Δ0 parameter estimates in female (dark orange) and male (light orange) rats. (D) Individual differences in the Δ+ parameter were predictive of oxycodone intake across female and male rats. (E) Individual differences in the Δ+ parameter were predictive of the A parameter in female and male rats. (F) Mediation analysis of sex, the Δ+ parameter, and initial strength of oxycodone reinforcement (e.g., A parameter).

To investigate the computational mechanism(s) underlying the differences in PRL performance, the parameter estimates obtained from the differential forgetting reinforcement learning algorithm were compared between the sexes. Post-hoc analyses of the significant sex-by-parameter interaction (Wald x^2^=24.50; p<0.001) indicated that the y_C_ and y_U_ were larger (y_C_: Wald x^2^=16.42; p<0.001; y_U_: Wald x^2^=7.27; p=0.007) and, notably, that the Δ_+_ parameter was lower (Wald x^2^=21.51; p<0.001) in female rats compared to male rats. A multiple regression analysis indicated that the Δ_+_ parameter explained the greatest amount of variation in PRL performance (Δ_+_ parameter: t=5.57; p<0.001, R^2^=0.37) across sexes (sex-by-Δ_+_ parameter interaction: t=1.79; p=0.08) compared to the other parameter estimates. The Δ_+_ parameter, however, did not fully account for sex differences in PRL performance (inclusion of sex in the regression model: R^2^ change=0.06; F_(1,112)_=10.67; p=0.001) suggesting that additional computational processes not directly measured here likely differ between male and female rats.

Nevertheless, these findings suggest that pre-existing disruptions in reward-guided decision-making may be the mechanism leading to sex differences in oxycodone self-administration. To directly test this hypothesis, sex and the Δ_+_ parameter were entered into a multiple regression analysis predicting total oxycodone intake. Sex was no longer a significant predictor of total oxycodone intake when the Δ_+_ parameter was included in the regression model (sex: t=0.31, p=0.76; Δ_+_ parameter: t=3.74; p<0.001) indicating that the differences in oxycodone intake between male and female rats were almost completely accounted for by variation in the Δ_+_ parameter (Figure 6D). A similar effect was observed when sex and the Δ_+_ parameter were entered into a multiple regression analysis predicting the *A* parameter, or the initial strength of oxycodone reinforcement (sex: t=0.37, p=0.72; Δ_+_ parameter: t=3.25; p=0.002), but not the B parameter, or the rate of growth in oxycodone intake (sex: t=3.15, p=0.002; Δ_+_ parameter: t=0.14; p=0.89).

We then conducted a mediation analysis to determine if the effect of sex on oxycodone-taking behaviors was mediated by variation in the Δ_+_ parameter. The significant relationship between sex and the A parameter (/J=0.19; p=0.04) was attenuated when the Δ_+_ parameter variable was included in the model (/J=0.03; p=0.73).

To determine if this reduction in the effect of sex on the A parameter was significant, a bootstrap analysis was performed using the PROCESS macro (Hayes, 2018). The indirect effect was significant (95% CI: 0.33-1.31) indicating that there was significant reduction in the effect of sex on the A parameter when the Δ_+_ parameter was included in the regression model (Figure 6F).

## Discussion

The current study provides new evidence into the biobehavioral mechanisms that regulate oxycodone-taking behaviors. Using translationally relevant behavioral and computational approaches, we report that disruptions in reward-guided, decision-making functions are predictive of greater oxycodone self-administration and addiction-like behaviors in rats. Rats with poor performance in a PRL task were found to self-administered greater amounts of oxycodone compared to rats with better PRL performance. Moreover, we provide evidence that greater oxycodone-taking behaviors in female rats is driven by pre-existing decision-making deficits compared to male rats. These data, collectively, demonstrate the utility of decision-making phenotypes for interrogating biologically based differences in opiate-taking behaviors and use of computational phenotypes to assessing addiction susceptibility in drug-naive individuals.

### Deficits in reward-guided, decision-making predict greater oxycodone use

Not all individuals that use oxycodone develop an opiate use disorder and there is emerging evidence that some individuals are more likely than others to transition from opiate use to abuse compared to others (Cheatle et al., 2019; Levis et al., 2021; Kendler et al., 2023). Identifying these at-risk individuals prior to initiation of oxycodone use, however, has been challenging. The results of the current study indicate that computationally characterized decision-making functions could serve as a behavioral predictor of problematic oxycodone use. We report that rats with poor performance in the PRL task self-administered greater amounts of oxycodone, were more motivated to obtain an oxycodone reinforcer, slower to extinguish drug-taking response, and had greater cue-induced reinstatement compared to rats with better PRL performance. Moreover, decision-making deficits that predicted oxycodone use were specific to reward-mediated updating of action values: rats with a lower Δ_+_ parameter were less likely to repeat a rewarded action and self-administered more oxycodone compared to rats with a higher Δ_+_parameter. These findings with oxycodone are similar to what we have previously reported for methamphetamine (Groman et al., 2019b) and cocaine (Groman et al., 2020) suggesting that deficits in the Δ_+_ parameter are likely to represent an addiction susceptibility phenotype.

Our characterization of latent drug-taking phenotypes indicates that rats with a lower Δ_+_parameter self-administer greater amounts of oxycodone, regardless of drug dose (0.05 mg/kg/infusion or 0.1 mg/kg/infusion), vehicle (saccharin or absence of saccharin), or effort requirement (FR1 vs FR3; see Figure 4I, above). The reinforcing effects of oxycodone appear, therefore, to be attenuated in rats with a low Δ_+_ parameter which could encourage rats to consume greater amounts of oxycodone to achieve the same euphoric state of rats with a high Δ_+_ parameter. Alternatively, rats with a lower Δ_+_ parameter may be less sensitive to negative effects of drugs of abuse that regulate or constrain drug-taking behaviors (e.g., sedation, tachycardia, etc). Indeed, there is evidence that lower sensitivity to alcohol sedation, but greater self-reported rewarding effects of alcohol, predict alcohol use disorder symptoms in humans (King et al., 2014). Additional studies investigating the behavioral and computational processes that lead to differences in the Δ_+_ parameter will undoubtedly provide insights into addiction susceptibility mechanisms.

### Sex differences in decision-making are associated with oxycodone use

Self-administration of oxycodone was found to be greater in female rats compared to males and the power function indicated that both the initial strength of oxycodone reinforcement and the rate of growth in oxycodone use was greater in females compared to males. Similar sex differences have been observed by others using oral and intravenous oxycodone self-administration (Fulenwider et al., 2020; Kimbrough et al., 2020; Zanni et al., 2020). We hypothesized – based on the negative relationship between the Δ_+_parameter and the A parameter – that reward-guided decision-making assessed prior to oxycodone exposure would be attenuated in female rats and importantly account for sex differences in the A, but not the B, parameter.

Indeed, we provide direct evidence supporting this hypothesis and propose that sex-mediated disruptions in reward updating is the mechanism by which differences in drug use emerge between males and females. The decision-making phenotypes did not, however, account for the sex differences in the B parameter. This suggests that the greater escalation in oxycodone use observed in female rats may be linked to other addiction susceptibility mechanisms. For example, sex differences in the pharmacokinetics of oxycodone (Doyle et al., 2023) are likely to lead to differences in drug-induced adaptations that regulate oxycodone intake.

Poor decision-making observed in female rats in the current study was surprising given that our previous studies examining decision-making functions across adolescent development in the rat have not observed decision-making differences between males and females (Moin Afshar et al., 2020; Villiamma et al., 2022). Sex differences in decision-making might, therefore, emerge during early adulthood when prefrontal circuits have stabilized (Casey et al., 2000; Galvan et al., 2006). Neuroimaging studies in humans have reported that prefrontal neurodevelopment plateaus earlier in females compared males (Lenroot et al., 2007; Heitzeg et al., 2018) which may lead to a precocial stabilization of decision-making functions in females. Prefrontal circuit stabilization does not depend on the presence or absence of gonadal hormones in females (Boivin et al., 2018) but may involve a more complex interaction between stress hormones, pubertal hormone exposure, and age (Galván and Rahdar, 2013; Spear, 2013). Additional longitudinal studies that quantify these factors of interest across the lifespan of individual are needed to understand the long-term impact of hormones on development and cognition.

### Neurobiology of reward-guided decision making

The neurobiology that regulates the Δ_+_ parameter is not known, but previous studies from our group have indicated a key role of midbrain dopamine signaling. We have previously reported that midbrain D3 receptor availability is negatively related to the Δ_+_parameter (Groman et al., 2016) and, notably, predictive of cocaine-taking behaviors (Groman et al., 2020). Midbrain D3 receptors act as autoreceptors and are likely to regulate the release of dopamine in brain regions known to be involved in decision-making, such as the amygdala and orbitofrontal cortex (Groman et al., 2019a). Individuals with greater midbrain D3 receptors may have lower phasic dopamine release particularly in response to rewards which leads to poorer value updating and, therefore, a lower Δ_+_ parameter. Indeed, a hypodopaminergic state has been proposed to be a risk factor for addiction (Goutaudier et al., 2022; Kótyuk et al., 2022) and associated with poor decision making (Rutledge et al., 2009, 2015; Clarke et al., 2011; Costa et al., 2015). Midbrain D3 receptors may, therefore, be at the nexus between addiction susceptibility and decision-making.

### Summary

By integrating sophisticated behavioral assessments with computational tools and self-administration assays, the current study provides new evidence indicating that poor reward-guided, decision-making is a predictor of problematic oxycodone use. Our data in female rats reveals a novel biobehavioral understanding of how differences in drug-taking behaviors might emerge between male and female subjects. Future studies integrating our computational approaches with drug-taking behaviors, neuroimaging, and genomic assays will provide critical insights into the neurobiological regulators of opiate use disorder.

## Acknowledgements and Disclosures

This work was supported by the National Institute on Drug Abuse (Grants DA051598 and DA051977). SMG was responsible for the conceptualization. SMG, PV, KL, JH, and MR were responsible for investigation. SMG was responsible for analysis and writing the original draft. SMG, PV, KL, JH, MR, and RJD approved the final version of the manuscript.

The authors report no biomedical financial interests or potential conflicts of interest.

## Notes

### Competing Interest Statement

The authors have declared no competing interest.

## References

Barraclough DJ, Conroy ML, Lee D (2004) Prefrontal cortex and decision making in a mixed-strategy game. Nat Neurosci 7:404–410 Available at: http://www.ncbi.nlm.nih.gov/pubmed/15004564.

Boivin JR, Piekarski DJ, Thomas AW, Wilbrecht L (2018) Adolescent pruning and stabilization of dendritic spines on cortical layer 5 pyramidal neurons do not depend on gonadal hormones. Dev Cogn Neurosci 30:100–107.

Carrasco-Garrido P, Gallardo-Pino C, Jiménez-Trujillo I, Hernández-Barrera V, García-Gómez-Heras S, Lima Florencio L, Palacios-Ceña D (2022) Nationwide Population-Based Study About Patterns of Prescription Opioid Use and Misuse Among Young Adults in Spain. Int J Public Health 67 Available at: /pmc/articles/PMC9437214/ [Accessed March 8, 2024].

Casey BJ, Giedd JN, Thomas KM (2000) Structural and functional brain development and its relation to cognitive development. Biol Psychol 54:241–257 Available at: https://www.sciencedirect.com/science/article/pii/S0301051100000582 [Accessed March 21, 2018].

Cheatle MD, Compton PA, Dhingra L, Wasser TE, O’Brien CP (2019) Development of the Revised Opioid Risk Tool to Predict Opioid Use Disorder in Patients with Chronic Nonmalignant Pain. J Pain 20:842–851.

Clarke HF, Hill GJ, Robbins TW, Roberts AC (2011) Dopamine, but not serotonin, regulates reversal learning in the marmoset caudate nucleus. J Neurosci 31:4290– 4297 Available at: http://www.ncbi.nlm.nih.gov/pubmed/21411670.

Costa VD, Tran VL, Turchi J, Averbeck BB (2015) Reversal Learning and Dopamine: A Bayesian Perspective. J Neurosci 35:2407–2416 Available at: http://www.ncbi.nlm.nih.gov/pubmed/25673835 [Accessed July 3, 2017].

Doyle MR, Martinez AR, Qiao R, Dirik S, Di Ottavio F, Pascasio G, Martin-Fardon R, Benner C, George O, Telese F, de Guglielmo G (2023) Strain and sex-related behavioral variability of oxycodone dependence in rats. Neuropharmacology 237:109635.

Enga RM, Jackson A, Damaj MI, Beardsley PM (2016) Oxycodone physical dependence and its oral self-administration in C57BL/6J mice. Eur J Pharmacol 789:75–80.

Evans CJ, Cahill CM (2016) Neurobiology of opioid dependence in creating addiction vulnerability. F1000Research 5 Available at: /pmc/articles/PMC4955026/ [Accessed March 8, 2024].

Evans WN, Lieber E, Power P (2018) How the Reformulation of OxyContin Ignited the Heroin Epidemic. Available at: https://www.nber.org/papers/w24475 [Accessed March 8, 2024].

Fulenwider HD, Nennig SE, Hafeez H, Price ME, Baruffaldi F, Pravetoni M, Cheng K, Rice KC, Manvich DF, Schank JR (2020) Sex differences in oral oxycodone self-administration and stress-primed reinstatement in rats. Addict Biol 25:e12822 Available at: https://onlinelibrary.wiley.com/doi/full/10.1111/adb.12822 [Accessed March 8, 2024].

Galvan A, Hare TA, Parra CE, Penn J, Voss H, Glover G, Casey BJ (2006) Earlier development of the accumbens relative to orbitofrontal cortex might underlie risk-taking behavior in adolescents. J Neurosci 26:6885–6892.

Galván A, Rahdar A (2013) The neurobiological effects of stress on adolescent decision making. Neuroscience 249:223–231.

Goutaudier R, Joly F, Mallet D, Bartolomucci M, Guicherd D, Carcenac C, Vossier F, Dufourd T, Boulet S, Deransart C, Chovelon B, Carnicella S (2022) Hypodopaminergic state of the nigrostriatal pathway drives compulsive alcohol use. Mol Psychiatry 2022 281 28:463–474 Available at: https://www.nature.com/articles/s41380-022-01848-5 [Accessed March 25, 2024].

Groman S, Hillmer A, Liu H, Fowles K, Holden D, Morris E, Lee D, Taylor J (2020) Midbrain D 3 Receptor Availability Predicts Escalation in Cocaine Self-administration. Biol Psychiatry 88:767–776 Available at: https://pubmed.ncbi.nlm.nih.gov/32312578/ [Accessed November 4, 2021].

Groman SM, Jentsch JD (2012) Cognitive control and the dopamine D 2-like receptor: A dimensional understanding of addiction. Depress Anxiety 29.

Groman SM, Keistler C, Keip AJ, Hammarlund E, DiLeone RJ, Pittenger C, Lee D, Taylor JR (2019a) Orbitofrontal Circuits Control Multiple Reinforcement-Learning Processes. Neuron 103:734–746.e3 Available at: http://www.ncbi.nlm.nih.gov/pubmed/31253468 [Accessed August 23, 2019].

Groman SM, Lee B, London ED, Mandelkern MA, James AS, Feiler K, Rivera R, Dahlbom M, Sossi V, Vandervoort E, Jentsch JD (2011) Dorsal striatal D2-like receptor availability covaries with sensitivity to positive reinforcement during discrimination learning. J Neurosci 31:7291–7299.

Groman SM, Massi B, Mathias SR, Lee D, Taylor JR (2019b) Model-Free and Model-Based Influences in Addiction-Related Behaviors. Biol Psychiatry 85:936–945

Available at: https://linkinghub.elsevier.com/retrieve/pii/S0006322318321218 [Accessed September 21, 2019].

Groman SM, Smith NJ, Petrullli JR, Massi B, Chen L, Ropchan J, Huang Y, Lee D, Morris ED, Taylor JR (2016) Dopamine D3 Receptor Availability Is Associated with Inflexible Decision Making. J Neurosci 36:6732–6741 Available at: http://www.ncbi.nlm.nih.gov/pubmed/27335404 [Accessed December 20, 2019].

Groman SM, Thompson SL, Lee D, Taylor JR (2022) Reinforcement learning detuned in addiction: integrative and translational approaches. Trends Neurosci 45:96–105.

Han B, Compton WM, Blanco C, Crane E, Lee J, Jones CM (2017) Prescription opioid use, misuse, and use disorders in U.S. Adults: 2015 national survey on drug use and health. Ann Intern Med 167:293–301 Available at: https://annals.org [Accessed March 8, 2024].

Hayes AF (2018) Introduction to Mediation, Moderation, and Conditional Process Analysis, Second Edition: A Regression-Based Approach. Guilford Press 46:1–692 Available at: https://www.guilford.com/books/Introduction-to-Mediation-Moderation-and-Conditional-Process-Analysis/Andrew-Hayes/9781462534654 [Accessed November 22, 2021].

Heitzeg MM, Hardee JE, Beltz AM (2018) Sex differences in the developmental neuroscience of adolescent substance use risk. Curr Opin Behav Sci 23:21–26 Available at: /pmc/articles/PMC6028191/ [Accessed March 25, 2024].

Ito M, Doya K (2009) Validation of decision-making models and analysis of decision variables in the rat basal ganglia. J Neurosci 29:9861–9874 Available at: http://www.ncbi.nlm.nih.gov/pubmed/19657038.

Kenan K, Mack K, Paulozzi L (2012) Trends in prescriptions for oxycodone and other commonly used opioids in the United States, 2000–2010. Open Med 6:e41 Available at: /pmc/articles/PMC3659213/ [Accessed March 8, 2024].

Kendler KS, Lönn SL, Ektor-Andersen J, Sundquist J, Sundquist K (2023) Risk factors for the development of opioid use disorder after first opioid prescription: a Swedish national study. Psychol Med 53:6223–6231 Available at: https://www.cambridge.org/core/journals/psychological-medicine/article/risk-factors-for-the-development-of-opioid-use-disorder-after-first-opioid-prescription-a-swedish-national-study/5AC2B3A018FD02073B5AD618FB741696 [Accessed March 25, 2024].

Kibaly C, Alderete JA, Liu SH, Nasef HS, Law PY, Evans CJ, Cahill CM (2020) Oxycodone in the Opioid Epidemic: High ‘Liking’, ‘Wanting’, and Abuse Liability. Cell Mol Neurobiol 2020 415 41:899–926 Available at: https://link.springer.com/article/10.1007/s10571-020-01013-y [Accessed March 8, 2024].

Kimbrough A, Kononoff J, Simpson S, Kallupi M, Sedighim S, Palomino K, Conlisk D, Momper JD, de Guglielmo G, George O (2020) Oxycodone self-administration and withdrawal behaviors in male and female Wistar rats. Psychopharmacology (Berl) 237:1545–1555 Available at: https://pubmed.ncbi.nlm.nih.gov/32114633/ [Accessed March 25, 2024].

King AC, McNamara PJ, Hasin DS, Cao D (2014) Alcohol Challenge Responses Predict Future Alcohol Use Disorder Symptoms: A 6-Year Prospective Study. Biol Psychiatry 75:798 Available at: /pmc/articles/PMC4280017/ [Accessed March 20, 2024].

Kótyuk E, Potenza MN, Blum K, Demetrovics Z (2022) The Reward Deficiency Syndrome and Links with Addictive and Related Behaviors. Handb Subst Misuse Addict From Biol to Public Heal:59–74 Available at: https://link.springer.com/referenceworkentry/10.1007/978-3-030-92392-1_3 [Accessed March 25, 2024].

Lankenau SE, Teti M, Silva K, Bloom JJ, Harocopos A, Treese M (2012) Initiation into Prescription Opioid Misuse among Young Injection Drug Users. Int J Drug Policy 23:37 Available at: /pmc/articles/PMC3196821/ [Accessed September 28, 2023].

Lenroot RK, Gogtay N, Greenstein DK, Wells EM, Wallace GL, Clasen LS, Blumenthal JD, Lerch J, Zijdenbos AP, Evans AC, Thompson PM, Giedd JN (2007) Sexual dimorphism of brain developmental trajectories during childhood and adolescence. Neuroimage 36:1065–1073.

Levis SC, Mahler S V., Baram TZ (2021) The Developmental Origins of Opioid Use Disorder and Its Comorbidities. Front Hum Neurosci 15:601905.

Marie N, Noble F (2023) Oxycodone, an opioid like the others? Front Psychiatry 14:1229439 Available at: https://www.molinspiration.com [Accessed March 8, 2024].

Moin Afshar N, Cinotti F, Martin DA, Khamassi M, Calu DJ, Taylor JR, Groman SM (2022) Reward-mediated, model-free reinforcement-learning mechanisms in Pavlovian and instrumental tasks are related. J Neurosci:JN-RM-1113-22 Available at: https://pubmed.ncbi.nlm.nih.gov/36216504/ [Accessed November 7, 2022].

Moin Afshar N, Keip AJ, Taylor JR, Lee D, Groman SM (2020) Reinforcement learning during adolescence in rats. J Neurosci 40:5857–5870. Available at: http://www.jneurosci.org/lookup/doi/10.1523/JNEUROSCI.0910-20.2020 [Accessed July 27, 2020].

Monico LB, Mitchell SG (2018) Patient perspectives of transitioning from prescription opioids to heroin and the role of route of administration. Subst Abus Treat Prev Policy 13:1–8 Available at: https://substanceabusepolicy.biomedcentral.com/articles/10.1186/s13011-017-0137-y [Accessed March 8, 2024].

Moningka H, Lichenstein S, Yip SW (2019) Current Understanding of the Neurobiology of Opioid Use Disorder: an Overview. Curr Behav Neurosci Reports 6:1–11 Available at: https://link.springer.com/article/10.1007/s40473-019-0170-4 [Accessed March 8, 2024].

Oswald LM, Dunn KE, Seminowicz DA, Storr CL (2021) Early Life Stress and Risks for Opioid Misuse: Review of Data Supporting Neurobiological Underpinnings. J Pers Med 2021, Vol 11, Page 315 11:315 Available at: https://www.mdpi.com/2075-4426/11/4/315/htm [Accessed March 8, 2024].

Payer DE, Behzadi A, Kish SJ, Houle S, Wilson AA, Rusjan PM, Tong J, Selby P, George TP, McCluskey T, Boileau I (2014) Heightened D(3) dopamine receptor levels in cocaine dependence and contributions to the addiction behavioral phenotype: a positron emission tomography study with [(11)C]-(+)-PHNO. Neuropsychopharmacology 39:311–318 Available at: http://www.ncbi.nlm.nih.gov/pubmed/23921256.

Richesson D, Jennifer M. Hoenig (2021) Key Substance Use and Mental Health Indicators in the United States: Results from the 2020 National Survey on Drug Use and Health.

Rutledge RB, Lazzaro SC, Lau B, Myers CE, Gluck MA, Glimcher PW (2009) Dopaminergic drugs modulate learning rates and perseveration in Parkinson’s patients in a dynamic foraging task. J Neurosci 29:15104–15114 Available at: http://www.ncbi.nlm.nih.gov/pubmed/19955362.

Rutledge RB, Skandali N, Dayan P, Dolan RJ (2015) Dopaminergic Modulation of Decision Making and Subjective Well-Being. J Neurosci 35:9811–9822 Available at: https://www.jneurosci.org/content/35/27/9811 [Accessed March 25, 2024].

Slivicki RA, Earnest T, Chang YH, Pareta R, Casey E, Li JN, Tooley J, Abiraman K, Vachez YM, Wolf DK, Sackey JT, Kumar Pitchai D, Moore T, Gereau RW, Copits BA, Kravitz A V., Creed MC (2023) Oral oxycodone self-administration leads to features of opioid misuse in male and female mice. Addict Biol 28:e13253 Available at: https://onlinelibrary.wiley.com/doi/full/10.1111/adb.13253 [Accessed March 8, 2024].

Spear LP (2013) Adolescent neurodevelopment. J Adolesc Heal 52:S7.

Villiamma P, Casby J, Groman SM (2022) Adolescent reinforcement-learning trajectories predict cocaine-taking behaviors in adult male and female rats. Psychopharmacology (Berl) 239:2885–2901 Available at: https://pubmed.ncbi.nlm.nih.gov/35705734/ [Accessed March 6, 2023].

Volkow ND, Fowler JS, Wang GJ, Hitzemann R, Logan J, Schlyer DJ, Dewey SL, Wolf AP (1993) Decreased dopamine D2 receptor availability is associated with reduced frontal metabolism in cocaine abusers. Synapse 14:169–177 Available at: http://www.ncbi.nlm.nih.gov/entrez/query.fcgi?cmd=Retrieve&db=PubMed&dopt=Citation&list_uids=8101394.

Webster LR (2017) Risk Factors for Opioid-Use Disorder and Overdose. Anesth Analg 125:1741–1748 Available at: https://pubmed.ncbi.nlm.nih.gov/29049118/ [Accessed March 8, 2024].

Zanni G, DeSalle MJ, Deutsch HM, Barr GA, Eisch AJ (2020) Female and male rats readily consume and prefer oxycodone to water in a chronic, continuous access, two-bottle oral voluntary paradigm. Neuropharmacology 167:107978.

